# Customizable, engineered substrates for rapid screening of cellular cues

**DOI:** 10.1101/693598

**Authors:** Eline Huethorst, Marie FA Cutiongco, Fraser A Campbell, Anwer Saeed, Rachel Love, Paul M Reynolds, Matthew J Dalby, Nikolaj Gadegaard

## Abstract

Biophysical cues robustly direct cell responses and are thus important tools for *in vitro* and translational biomedical applications. High throughput platforms exploring substrates with varying physical properties are therefore valuable, however, currently existing platforms are limited in throughput, the biomaterials used, the capability to segregate between different cues and the assessment of dynamic cellular responses. Here we present a multiwell array (3×8) using a substrate engineered with patterns that present topography or rigidity cues welded to a bottomless plate with a 96-well format. Both the patterns on the engineered substrate and the well plate format can be easily customized, permitting systematic and efficient screening of biophysical cues. Here, we demonstrate three multiwell arrays patterned with a variety of topographical and mechanical cues (nano-grooves, soft pillars and nano pillars) tested with three different cell types. Using the multiwell array, we were able to measure cell functionality using analytical modalities such as live microscopy, qPCR and fluorescent immunochemistry. Cardiomyocytes cultured on 5µm grooves showed less variation in electrophysiology and contractile function. Nanopillars with 127 nm height, 100 nm diameter and 300 nm pitch showed improved chondrogenic maintenance from matrix deposition and chondrogenic gene expression. High aspect ratio pillars with an elastic shear modulus of 16 kPa mimicking the cortical bone altered cell adhesion, morphology, and increased expression of osteogenic genes. We have demonstrated the bespoke, controlled and high-throughput properties of the multiwell array that are currently unparalleled in the field today.

---

Through its ability to regulate cell behavior, the cellular micro-environment plays a key role in health and disease.^1-4^ Control of cell behavior using appropriate environmental cues is therefore important to understand as *in vitro* tools that aid experimentation and that can guide directed repair and regeneration of tissues *in vivo are being sought.*^5,6^ Reproduction of the biochemical and biophysical microenvironment can be achieved in a precise and reproducible manner through controlled nanoscale materials.^7-10^ Of particular interest are engineered substrates with defined biophysical properties that can recapitulate the substrate rigidity or surface topographical cues present in the natural cell environment.^11,12^ Precisely engineered topographical cues or substrate rigidities have been shown to preferentially direct mesenchymal stem cell differentiation,^13-15^ alter endothelial cell functionality^16-18^ and change in neurogenic subtype.^19,20^

Testing of engineered substrates has long relied on the use of individual substrates assessed in tandem to screen for positive hits but is severely hindered in throughput. In recent years, combinatorial libraries of biomaterials, including topographies, have been made to increase efficiency of screening.^10,21-23^ However, these current high-content platforms lack physical segregation between substrates of interest. Continuous exchange of signaling molecules between cells makes it impossible to uncouple biophysical and paracrine based effects. In addition, these combinatorial libraries have bespoke dimensions incompatible with most analytical laboratory equipment. New platforms that allow rapid and high-throughput screening of a library of materials are thus required. A good screening platform should also be able to isolate the effect of a specific biophysical cue to limit confounding paracrine effects^24,25^ and should be made of biocompatible materials. Moreover, these screening platforms need to be highly generalizable across substrates, cell types and various regenerative medicine applications. The screening platform should additionally allow a wide variety of validation assays for thorough selection of the most appropriate features for possible translational application.

In this study, we present a new platform for rapid screening of nanotopographies with altered biophysical properties. The multiwell array is a robust and high throughput platform based on thermoplastics such as polystyrene, with the footprint and dimensions of a 96-well plate. The complete multiwell array is a fully customizable slide welded to a bottomless well plate, both of which were manufactured through injection moulding. This allows for an industrial level production of biocompatible substrates with low cost and high reproducibility. The multiwell array is presented in a 96-well format, allowing various biological assays to be carried out with standard laboratory tools and techniques. This includes quantitative polymerase chain reaction (qPCR), fluorescent immunochemistry, immunohistochemistry and microscopy. With this design, 24 different topographies or rigidities can be simultaneously compared without any paracrine signals influencing results. We showed that the multiwell array can be customized to contain a wide array of mechanical or topographical cues in a high-throughput fashion, allowing us to assess behavioral changes of various cell types within the same platform. Thus, the multiwell array presents an alternative screening platform for rapid, accurate and highly reproducible interrogation of new engineered microenvironments.

## Results

### Customization of the multiwell array

The multiwell array is comprised of two parts, each fully customizable in design. An overview of the fabrication process is depicted in **Figure 1** and detailed in the **Experimental Section**. First, topographies or rigidities of interest (defined by patterns) are created on a slide through a multistep engineering process. A master stamp containing the patterns of interest are defined on silicon (Figure 1A) or quartz (Figure 1B) through standard fabrication techniques of electron beam lithography (EBL) and plasma etching. The master stamps are customized by combining different shapes (*i.e.* pits, pillars and grooves) and length scales from nano- to micrometer sizes depending on the specifications of interest.

**Figure 1.**
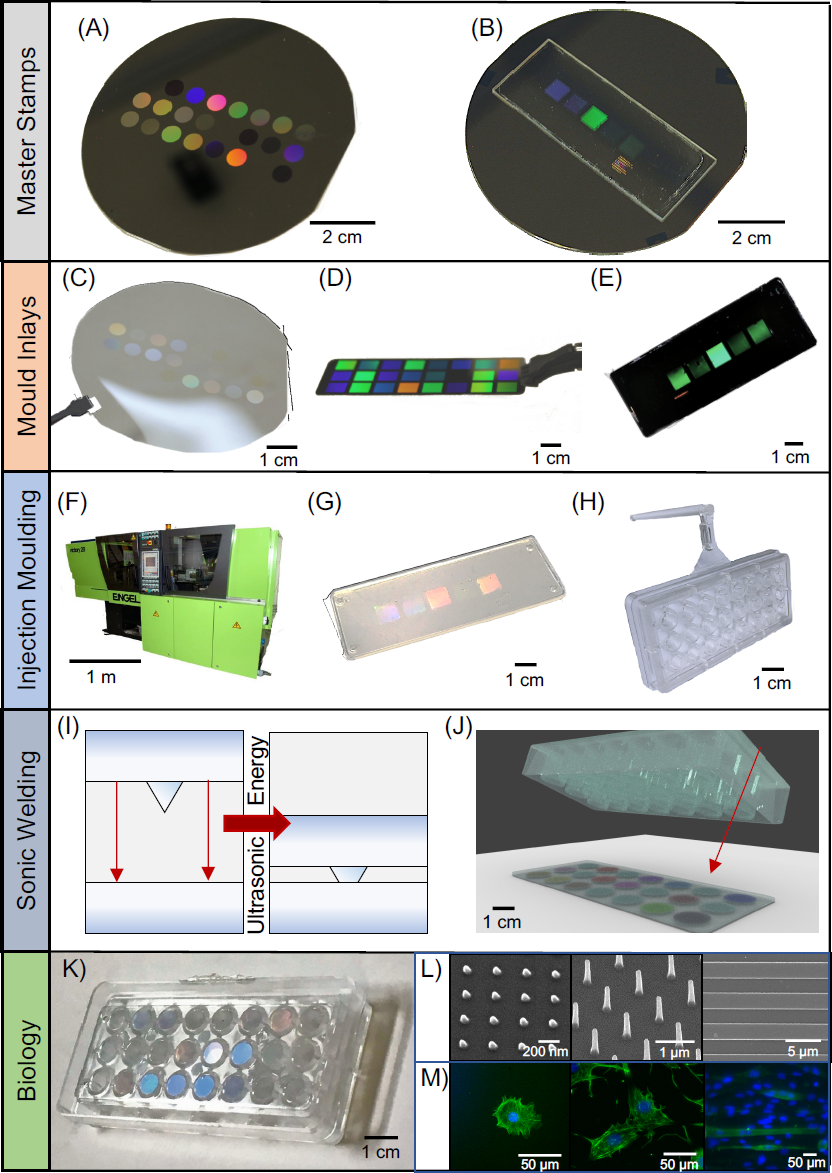
Bespoke multiwell array fabrication for rapid screening of rigidity or topographical cues. We illustrate the various stages involved in the fabrication of our highly customizable multiwell array. See Experimental Section for the detailed processes. First, master stamps are fabricated with the desired patterns on (A) a silicon wafer or (B) a quartz slide. The initial pattern is formed using electron beam lithography (EBL) and metal lift-off, and then etched into features using a plasma etching process. Pillars, pits or grooves can be defined on master stamps to fabricate multiwell arrays that provide topographical cues. High aspect ratio pillars can be defined on master stamps to provide controlled changes in substrate rigidity. Afterwards, a negative relief of the master stamp is fabricated using (C) nanoimprint lithography (NIL) to make a SmartNIL (EVG) foil, (D) electroplating of nickel or (E) NIL of SU-8 epoxy photoresist on Cirlex® polyimide.^28^ The resulting polymer replica are used in an (F) industrial grade injection moulding Engel Victory tool, which (G) moulds thermoplastic polymers such as polystyrene or polycarbonate to replicate structures of the original master stamp onto a slide. (H) A bottomless well plate with 8 columns x 3 row and approximately 0.3cm^2^ growth area (similar to standard 96 well plate) was also made from injection moulding of polystyrene. (I-J) To unite the slide and the bottomless well plate, the two are brought into contact and ultrasonic energy is used to melt a weld seam on the plate into the replica to form a joint around the patterned area. (K) The multiwell array combines multiple types of biophysical cues in one plate, (L) *e.g.* nano-pillars, tall pillars or nano- and micro-grooves, and (M) multiple cell types. This allows for high-throughput screening of individual cues without risk of paracrine signalling confounding the effects of topography or rigidity. The standard well plate format of the multiwell array allows established analytical techniques such as microscopy to be performed easily.

To enable high-throughput production of the engineered substrate, the master stamps are then used to create negative relief replica to be used as mould inlay for injection moulding (**Figure 1C-E**). The mould inlay is either in nickel or polymeric material to withstand high temperatures and high pressures required for high fidelity replication of patterns using injection moulding^26^ (**Figure 1F**). From one mould inlay, hundreds of slides containing the patterns as the original master stamp per hour are made through injection moulding (**Figure 1G**). We have not seen deterioration of the mould inlay after hundreds to thousands of replicates. But when needed, the original master stamp can be used again to create a new mould inlay or a new master stamp fabricated for further production of slides.

Aside from the engineered substrate slide containing patterns, the well plate format can be easily tuned to match the scale of the experiment required by the end user. The bottomless well plate is also produced through the same high throughput injection moulding process. The dimensions and arrangement of the patterns on the master stamp are set to match the specifications of the desired well plate format. In this study, we focus on creating a multiwell array, containing 24 wells with a 96-well format (0.3 cm^2^ per well), which is one of the most commonly used and preferred formats for automated and high throughput screening (**Figure 1H**).^27^

For a fully enclosed device, the two components are joined together through ultrasonic welding (**Figure I-J**). Since the slide and bottomless well plate are made separately, one can mix and match different combinations of the two components easily. Here, we demonstrate how we created multiwell arrays that presented different nanopillars, grooves or high aspect ratio pillars all in the same 96-well format. These topographies and rigidity cues incorporated in the multiwell array were then used to test changes in functionality of different cell types. To show the utility of our customized multiwell array we developed polystyrene and polycarbonate slides patterned with varying topographies and rigidities and screened them on the behavior of cardiomyocytes, chondrocytes and osteoblasts.

### Multiwell array for physiological real-time assessment of hiPSC-CM function

Human induced pluripotent stem cell derived cardiomyocytes (hiPSC-CM) have been shown to elongate when cultured on microgrooves.^29^ More importantly, this elongation has been shown to improve functionality towards a more mature phenotype ^30-32^ As hiPSC-CMs exhibit a relatively immature phenotype compared to adult CMs, this strategy could be used to induce functional maturation of hiPSC-CM. We sought to isolate the interaction between hiPSC-CM and grooves from paracrine effects to understand the responses induced by groove topographies on cells. Thus, we screened grooves with various dimensions for hiPSC-CM morphology and functionality using a multiwell groove array. Because hiPSC-CM previously showed increased maturation on the most narrow features (8-30 µm wide),^1^ we chose to use a multiwell array with groove widths of 100 nm, 250 nm, 500 nm, 1000 nm, 2000 nm and 5000 nm and a width:pitch ratio of 1:1. The groove depth was kept constant at 250 nm (**Figure 2A**). We used a Flat surface as a control.

**Figure 2.**
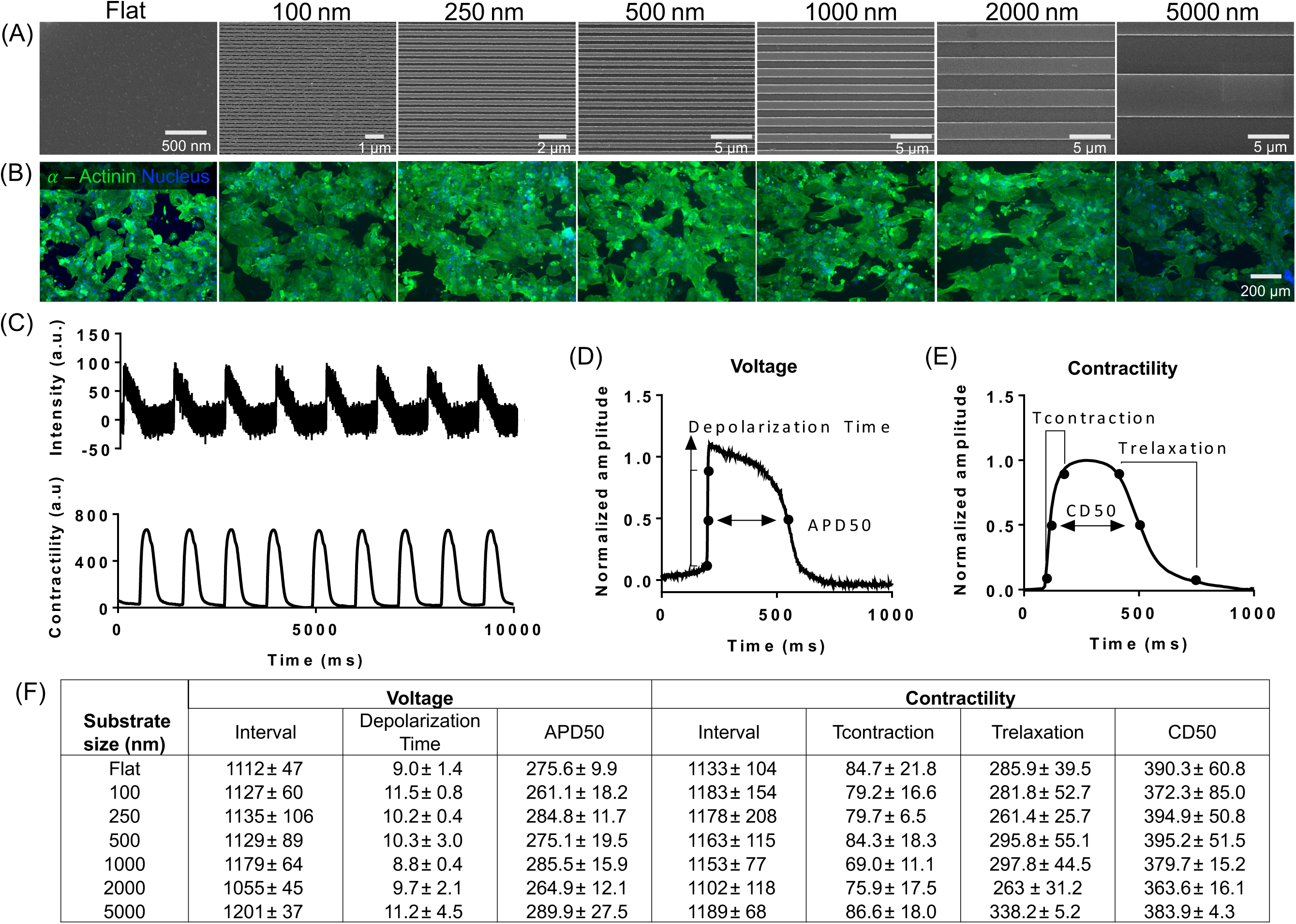
Functional stimulation of human induced pluripotent stem cell derived cardiomyocytes (hiPSC-CM) using grooves on the multiwell array. hiPSC-CM were grown on the grooves for 10 days before immunofluorescent staining and functional assessment using high speed microscopy. (A) Scanning electron micrographs (SEM) of grooves with widths of 100 nm, 250 nm, 500 nm, 1000 nm, 2000 nm or 5000 nm. All grooves had constant depth of 250 nm and width:pitch ratio at 1:1. (B) Fluorescent images of hiPSC-CM stained for α-actinin (green) and DAPI (blue) on corresponding groove topographies. (C) Functionality of hiPSC-CM were assessed by measuring voltage (at 10,000 frames per second (fps)) and contractile function (100 fps) for 10 s. Across 10 s, recordings were averaged from which various parameters were measured: (D) From averaged recordings of voltage, interval, depolarization time, and action potential duration at 50% of the amplitude (APD50) were measured. (E) From averaged recordings of contractility, contraction time (T_contraction_), relaxation time (T_relaxation_) and contraction duration at 50% of the amplitude (CD50) were measured. (F) Summary of functionality measurements of hiPSC-CM on different grooves. Data are shown as mean ± standard deviation (SD), with measurements obtained from n≥2 replicates. Statistical analysis using one-way ANOVA with a Dunnett post-hoc test showed no significant differences in functionality between groups.

While the nano- and micro-grooves do not affect morphology (**Figure 2B**) and electrophysiology (**Figure 2F**) of hiPSC-CMs after 10 days, the contractile behavior of hiPSC-CM is less varied (**Figure 2F**). More specifically, the CD50 and T_relaxation_ for this topography only had a SD of 4.3 ms and 5.2 ms, respectively, where all other substrates had SD ranging from 15.2 to 85.0 ms and 25.7 to 55.1 ms, respectively. HiPSC-CM and other iPSC derived cell types are known for their variability as a result of differences in donor and the protocols used for dedifferentiation.^33,34^ This is undesired as it has implications for experimental outcomes, including drug screens. Therefore, the reduction in well-to-well variability is very valuable. Nonetheless, more experimental repeats are needed to draw final conclusions. Example traces over time for intensity of voltage-sensitive dyes and contractile behavior measured from live brightfield microscopy (**Supplementary Videos**) are shown in **Figure 2C.** Parameter explanations for voltage and contractility are given in **Figures 2D and 2E**, respectively.

### Improved chondrogenic maintenance using nanopillars

Loss of chondrocyte phenotype and dedifferentiation into fibroblasts, commonly observed on standard tissue culture plastic, is exacerbated by increased adhesion to its substrate.^36^ We hypothesized that reduction of chondrocyte adhesion using nanopillars improves chondrocyte phenotype. A multiwell array with 14 nanopillar types (with fixed dimensions of 100 nm diameter and 300 nm pitch, and height varying from 27 nm to 205 nm) were used (**Figure 3A**). Using a variety of functional assays, we tested the effect of the nanopillar array in reversing dedifferentiation of chondrocytes previously cultured on tissue culture plastic (‘cultured chondrocytes’, **Figure 3B**). For visualization and ease of comparison, the data were presented in a heatmap. Over 28 days of culture, we observed that nanopillars with 62, 77, 127 and 190 nm heights changed chondrocyte behavior significantly. Compared to shorter nanopillars, cultured chondrocytes on tall nanopillars (height ≥ 127 nm) generally exhibited decreased proliferation and increased glycosaminoglycan deposition, indicating increased commitment of cells to the chondrocytic lineage.^37^ Chondrogenic function was also observed through gene expression analysis, where expression of SRY-Box 9 *(SOX9)* and collagen 2α1/collagen 1α1 (*COL2A1/COL1A1)* ratio was enhanced on nanopillars with 127 nm height compared to Flat. Expression of aggrecan (*ACAN*), a proteoglycan secreted by mature chondrocytes, and *SOX9* were also significantly upregulated by nanopillars with 190 nm height. On the other hand, chondrogenic genes *SOX9* and *COL2A1/COL1A1* ratio was minimized on 62 and 77 nm tall nanopillars after 28 days indicating fibroblastic phenotype.

**Figure 3.**
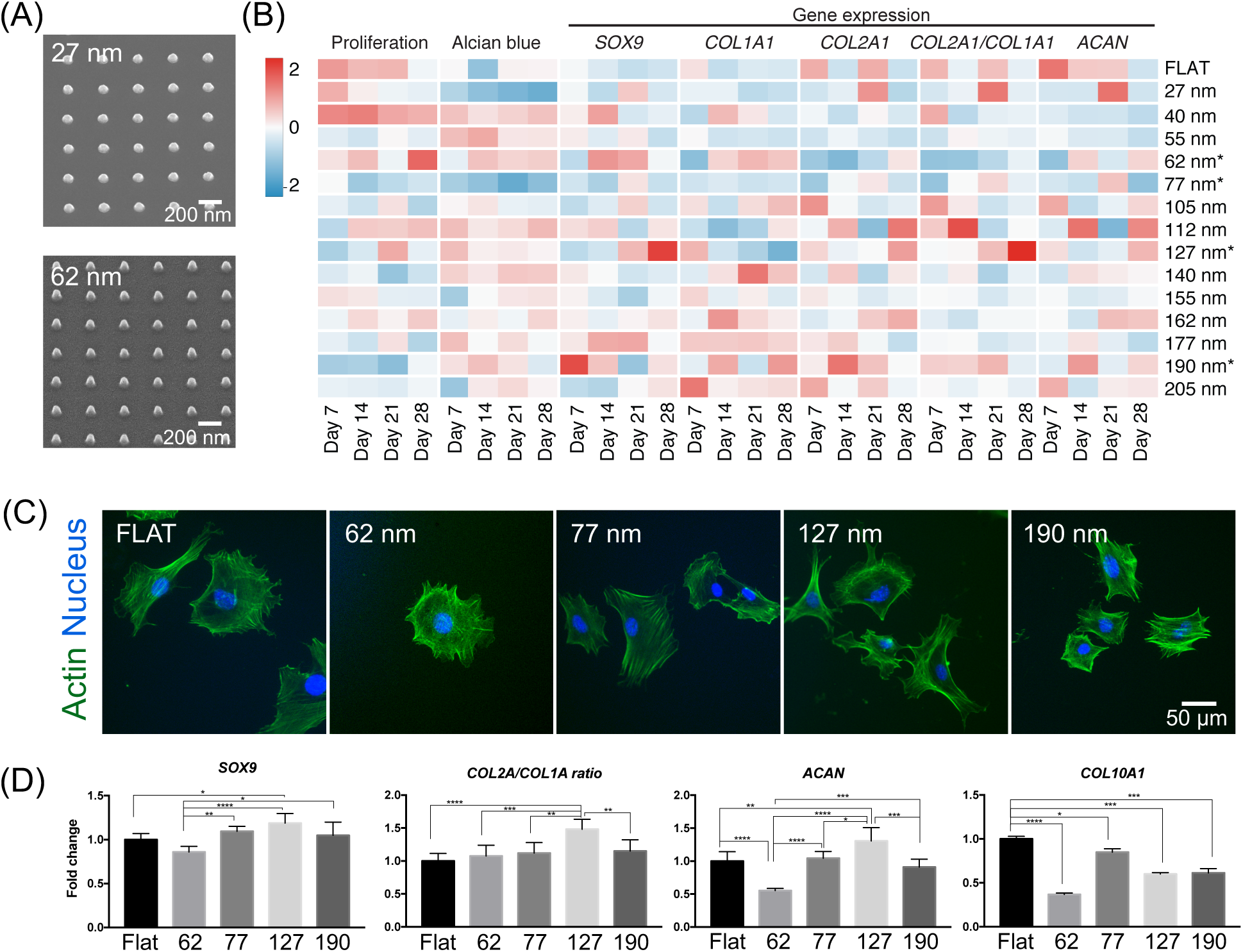
Tall nanopillars aid chondrogenic maintenance. A multiwell array containing nanopillars with constant 300 nm pitch, 100 nm diameter and varying heights were used to screen for topographies that improved or maintained chondrogenic properties of primary chondrocytes. Numbers denote the height of nanopillars examined. (A) SEM of representative nanopillars with heights of 27 and 62 nm found on the multiwell array. (B) Multimodal analysis of chondrogenic function induced by nanopillars over 28 days. A heatmap showing time point analysis of proliferation, matrix deposition and gene expression changed by nanopillars of varying height. Cultured chondrocytes were propagated in tissue culture plastic for 7 days before growth on the multiwell array to determine nanopillars that can reverse chondrogenic dedifferentiation. Chondrocyte behavior was measured at day 7 intervals for 28 days. Each tile represents the mean value of each measurement at a given time point across 2 independent experiments (n=4). Nanopillars that significantly changed behavior of cultured chondrocytes were selected for further examination. (C) Representative images of cultured chondrocytes on specific nanopillar heights after 24 hours. Cultured chondrocytes were fixed and stained against actin (green) and the nucleus (blue). (D) Expression of chondrogenic genes in primary chondrocytes after culture on selected nanopillars for 14 days. Gene expression data on chosen nanopillars are presented as mean ± SD across 3 independent experiments (n=6). Statistical significance was calculated using one-way ANOVA with Tukey’s post-hoc test with * denoting p <0.5, ** denoting p<0.05, *** denoting p <0.005, **** denoting p< 0.0001.

Cultured chondrocytes stained against the actin cytoskeleton also revealed changes in adhesion introduced by varying nanopillar heights (**Figure 3C**). Cultured chondrocytes on 62 and 77 nm heights generally were of larger size and more arranged actin stress fibers, similar to those observed on Flat. In contrast, taller nanopillars generally induced cells with more radially distributed actin fibers and smaller spread areas compared to Flat.

We then further selected nanopillars with 127 and 190 nm heights to improve the maintenance of freshly isolated primary (‘primary’) chondrocytes compared to standard tissue culture plastic (**Figure 3D**). We also included nanopillars with 62 and 77 nm heights as controls that were expected to deteriorate chondrogenic maintenance. Using primary chondrocytes we observed similar results as the cultured chondrocytes. At 14 days, *SOX9* and *COL2A1* and *ACAN* expression was significantly upregulated in primary chondrocytes by 127 nm high nanopillars compared with Flat. *COL1A1* expression was significantly reduced in 127 nm high nanopillars compared to Flat. Thus, the collagen *COL2A1*/*COL1A1* ratio in primary chondrocytes was significantly upregulated in 127 high nanopillars. However, all nanopillars reduced *COL10A1* expression in primary chondrocyte.

Using the multiwell nanopillar array we systematically screened for an optimal nanopillar height for chondrogenic differentiation and maintenance using a wide variety of standard analytical assays. Generally, we observed chondrogenic maintenance of primary isolated chondrocytes improved by nanopillars with 127 nm height compared to Flat. Nanopillars with 127 nm height represents a possible new material that could be used for sustained *in vitro* culture of primary chondrocytes without the need for expensive biochemical cues such as transforming growth factor (TGFβ).

### Substrate stiffness mimicked by nanopillars directs osteogenic differentiation

Osteogenic differentiation has been robustly shown to accelerate with higher substrate stiffness.^38^ Currently, no array format exists that allows for screening of cell response to tailored mechanical properties of the substrate. We fabricated a multiwell array with high aspect ratio nanopillars of varying diameter, pitch and height, to obtain surfaces that differ in stiffnesses (**Figure 4A**). Altering substrate stiffness by changing nanopillar dimensions is a highly controllable way of altering the rigidity compared to *e.g.* hydrogel stiffness that relies on tweaking chemical concentration or UV-light exposure. Furthermore, this extends the range of substrate rigidities available and removes complications of coupled biochemical and biophysical properties arising from chemically-defined biomaterials such as polyacrylamide.^39^ Cylindrical nanopillar arrays and bulk substrate mechanical properties have previously been demonstrated to be comparable using the effective shear modulus 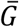, calculated as follows:^40^

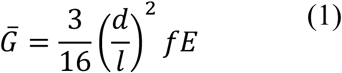

**Figure 4.**
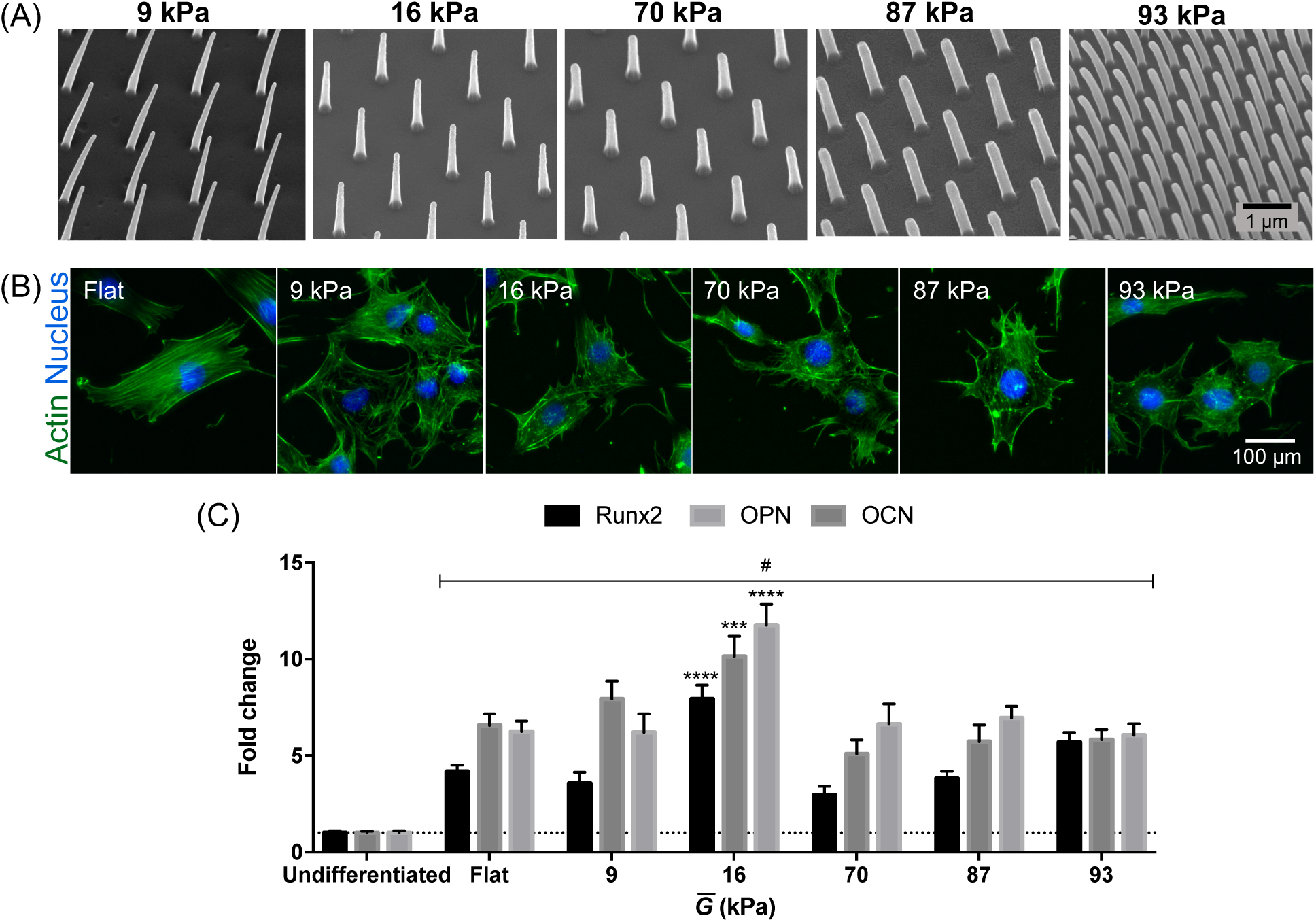
Substrate rigidity induces osteogenic differentiation. Substrate rigidity was controlled by varying the dimensions of high aspect ratio pillars. Substrate rigidity is reported as shear rigidity 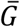. (A) SEM of high aspect ratio pillars with different 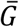 resulting from variations in pillar diameter and pitch. (B) Representative images of MC3T3 pre-osteoblast cells grown on varying rigidities after 24 hours. Cells were stained against actin (green) and nucleus (blue). Pre-osteoblast cells on varied substrate rigidities manifested drastic changes in morphology, especially size, filopodial formation and actin cytoskeleton organization compared to Flat. (C) Expression of osteogenic genes Runx family transcription factor 2 (*RUNX2)*, osteopontin (*OPN*) and osteocalcin (*OCN*) in MC3T3 pre-osteoblasts after 10 days of culture. # denotes statistically significant increase in osteogenic gene expression induced by all substrate rigidities for all genes compared to an Undifferentiated control (day 0). Fold change was calculated from f * denotes statistically significant increase in gene expression compared to Flat control. *** denotes p < 0.0005, **** denotes p < 0.0001. Statistical significance was measured using two-way ANOVA with Dunnett’s post hoc test. For comparison, Flat polycarbonate has shear modulus of 0.85 × 10^6^ kPa.

Where d is the diameter of the pillar tip, l is the length, or height, of the nanopillar, E is the Young’s modulus of the bulk material, and f is the fill factor of the array.

As the pillars demonstrated here differ in morphology from an ideal cylinder, the deflection characteristics of the pillars had to be established, and the effective shear modulus amended to account for this, which we will call 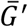. This was calculated using finite element analysis, and the discrepancy between the ideal cylinder and the modelled pillar calculated by comparing their spring constants, *Δk*. As this value is a constant, the amendment is simple:

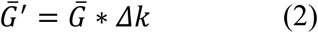

For comparison to bulk substrates, the shear modulus and the Young’s modulus are related by the Poisson’s ratio, providing that the substrate is, or can be treated as, isotropic:

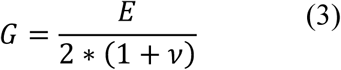

Where *ν* is Poisson’s ratio of the material. Dimensions of the high aspect ratio nanopillars and the corresponding mechanical properties are given in **Table S1**.

We cultured MC3T3 pre-osteoblast cells on high aspect ratio nanopillars without addition of biochemical inducers of osteogenesis. After 24 hours of culture, morphological differences in response to shear moduli were already manifested (**Figure 4B**). On Flat (with shear modulus of 0.85 × 10^6^ kPa), MC3T3 pre-osteoblasts showed spread morphology with organization of actin into highly aligned stress fibers highly characteristic of response on relatively stiff substrates. However, pre-osteoblast cells on varying substrate rigidities showed higher formation of filopodia and less organized actin structures. In particular, cells on 9 and 16 kPa rigidities induced formation of cortical actin and showed cells with circular shapes. Cells on 70 kPa and 87 kPa exhibited particularly long filopodial extensions but less circular and more elongated cell shapes compared to softer substrates. Increasing the rigidity to 93 kPa induced actin organization similar to substrates with 70 and 87 kPa rigidity, but reduced filopodial extensions. After 10 days of culture, all substrate rigidities and Flat significantly increased expression of osteogenic genes Runx family transcription factor 2 (*RUNX2)*, osteopontin (*OPN*) and osteocalcin (*OCN*) compared to an Undifferentiated control (**Figure 4C**). This was unsurprising as pre-osteoblasts tend to differentiate with increasing confluence.^41^ Though potent in changing MC3T3 pre-osteoblast morphology, substrate rigidities of 9, 70, 87 and 93 kPa showed similar osteogenic profile to a Flat control. Only the high aspect ratio nanopillars with 16 kPa shear modulus significantly increased expression for all three osteogenic markers compared to Flat. It has been reported that cortical bone has a Young’s modulus of 20-45 kPa.^38^ Applying equation (3) to this value and using a Poisson’s ratio of 0.3, gives a shear modulus of 8-17 kPa, which these high aspect ratio nanopillar arrays with 16 kPa shear modulus replicate. Here we observed that the stiffness of the bone microenvironment was effectively mimicked through simple manipulation of high aspect ratio pillars.

## Discussion and Conclusion

Finding the right engineered substrate to influence cell behavior is key for new *in vitro* tools and models of *in vivo* behavior, and for potential regenerative purposes. Without the ability to quickly engineer arrays of patterns specifying topographies or rigidities, discovery of positive hits remains limited and inefficient. Here, we fabricated a multiwell array that allows for multiple topographical or mechanical conditions to be assessed simultaneously. Tested against different cell types and using a variety of analytical techniques, we exhibited the flexibility and capability of the multiwell arrays for screening engineered substrates. The advantages of the multiwell array over currently available screening platforms^10,21,22^ are manifold, as discussed below.

First, the highly bespoke nature of the multiwell array enables creation of multitudes of mechanical and topographical cues to alter the cell microenvironment. Here, we have shown successful integration of a wide variety of patterns with different shapes (pillars vs grooves), length scales (nano- and micron-sized grooves), and effective rigidities (high aspect ratio pillars) in the multiwell array. Essentially, our method for multiwell array fabrication allows for any potential microenvironment exhibiting with geometric, topographical or mechanical properties to be mimicked with nanometer-scale precision. With our multiwell array method, both pattern replication throughput and fidelity, and cytocompatibility are improved. The current best patterned array available today utilizes the elastomer polydimethylsiloxane, which requires long curing times that ensure pattern replication but prevent high throughput production, may leak uncured oligomers toxic to cells,^42^ and provides a less reliable chemical interface.^43^

The multiwell array indeed lends large flexibility in configuration, allowing the end-user to explicitly make arrays specifically optimized for the task at hand. Aside from full customization in the patterned cues, the multiwell array can be scaled up to a full-sized well plate and customized to larger well plate formats. With larger arrays for cell growth, the multiwell array can be converted from a screening to an *in vitro* cell culture device. For instance, multiwell arrays formatted with 12-wells and patterned with 127 nm tall nanopillars could be manufactured as a new cell culture tool for improved chondrocyte maintenance. Multiwell arrays with high aspect ratio nanopillars of 16 kPa shear rigidity could be used as an alternative and low-cost method to stimulate osteogenesis compared to use of recombinant growth factors.

To elevate the variety of cell signals studied in the multiwell array, chemical cues may also be coated onto a flat polystyrene slide then welded to the bottomless well plate.^44,45^ Other thermoplastics (*e.g.* polyurethane^26^) or metal/thermoplastic and ceramic/thermoplastic composites^46^ amenable to forming complex microscale structures using injection moulding could be explored. The customizability of the mulitwell array truly permits screening of biophysical and biochemical environments.

Second, the multiwell array can be used for various analytical modes with standard laboratory equipment. One of the pitfalls of the currently available platforms is limitation of biological assessment to imaging techniques. Here, we showed that the multiwell array can be used with standard laboratory techniques such as fluorescent immunochemistry, qPCR, plate readers and microscopy. This allows for comprehensive examination of cell behavior induced by the engineered substrates on a genetic, morphological and functional level. We also exhibited that high-speed microscopic techniques for physiological measurements are possible using the multiwell array. Since containment is integrated with the patterned cues of interest, light only needs to with a single substrate. In terms of real-time techniques such as time lapse microscopy, the multiwell array also provides a stable platform that allows multiple locations to be assessed at once without regard to substrate drift. In contrast, standalone substrates (*e.g.* those made from soft lithography techniques) that require containment within a well will have two substrates interacting with light (i.e. the well plate and substrate on which cells adhere), and depending on density may be free floating in liquid.

Third, the multiwell array format provides patterned areas in isolation. Studies through time have shown that cells rapidly change their paracrine environment in response to the biochemical^47-49^ and biophysical^50^ milieu. In the multiwell array, we decouple the strong effects of paracrine factors freely available in the media (*e.g.* metabolites, cytokines and miRNAs) from confounding the mechano-transductive effects of the patterned cues in question.

Taken together, the multiwell array uniquely combines high-throughput production, flexibility in topographical and rigidity cues, and quality and customizability that no other screening platform to date offers.

## Experimental Section

### Nanopillars master stamp fabrication

A quartz substrate coated with a bilayer of poly(methyl methacrylate) (PMMA) was written with dots using electron beam lithography (EBL, Vistec VB6) using an ‘on the fly’ strategy as previously described^51^ After development in 1:1 methyl isobutyl ketone:isopropyl alcohol (MIBK:IPA) at 23°C, a 50 nm nichrome (NiCr) film was evaporated and lift-off performed in 50°C acetone for 12h. A sequence of five masked etches were then performed, alternating exposed patterns at each step and varying the etch depth. A positive-tone photoresist (Shipley S1818, microposit) was spun at 3000 rpm, exposed for 4.5s on a mask aligner (Suss MA6), and developed in 1:1 microposit developer:water for 75s. Nanopillar patterns were etched into the quartz substrate in a trifluoromethane / argon (CHF_3_/Ar) plasma in a reactive ion etching (RIE, Oxford RIE 80+) tool. Photoresist was removed in acetone, and the process was repeated with a different mask configuration until the 20 nanopillar patterns were etched to 20 different heights (5 iterations total). The slide was coated with a fluorosilane anti-stick layer, and an SU-8 epoxy photoresist / Cirlex polyimide (DuPont) hybrid inlay for injection moulding was patterned as a negative relief of the master using NIL as described previously.28

### Grooves master stamp fabrication

A silicon wafer was coated with a 200 nm film of a positive-tone resist (CSAR 62, AllResist) and patterns exposed using EBL with exposure time approximately 7h for 6.3cm^2^. Patterns were arranged in an 8 by 3 array on 9mm center-to-center pitch. Blank control regions were also included, and the pattern locations were randomized in the array. After EBL exposure, the wafer was developed in *n*-amyl acetate at 23°C and rinsed thoroughly in IPA. Grooves were transferred into the silicon substrate using sulfur hexafluoride / octafluorocyclobutane (SF_6_/C_4_F_8_) etching (STS inductively coupled plasma) to a depth of 250 nm. The remaining positive resist was removed in acetone. A NIL machine (EVG 5200) was used to create a polymer replica (SmartNIL foil) as a working stamp that was cut to size, mounted and used for injection moulding.

High aspect ratio nanopillars master stamp fabrication: A quartz slide was spincoated with a bilayer of PMMA, and the pattern was written using EBL. The pattern was developed for 1 min in 2.5:1 MIBK:IPA solution, and rinsed with IPA for 30 s. Residual PMMA in the nanopits was removed using a 30 s 80 W O_2_ plasma treatment, and an 80 mm thick layer of nickel was thereafter deposited. This was removed using the N-Methyl-2-pyrrolidone remove (remover 1165, microposit) at 50°C for 12 hours to form nanodots on nickel. These were then etched into nanopillar arrays using a CHF_3_/Ar plasma using reactive ion etching (Oxford RIE 80+) for 33 min in a single etch step process. Similar to the nanopillars array, a polymer replica was created from the nanopillar arrays through NIL of the SU-8 / Cirlex hybrid inlay.

### Injection moulding

Nanopillars, grooves and high aspect ratio pillars were injection moulded using polymer replica inserts mounted in a custom tooling configuration in an Engel Victory 28 injection moulder.^28,52^ The resulting polymer slides contained replica of structures found in the original master stamp (on either silicon or quartz) or negative relief of the polymer replica (on either smartNIL foil, nickel or SU-8/Cirlex polyimide). Nanopillars and grooves were injection mouleded in polystyrene (1810 crystal polystyrene, Total, Belgium), as previously described.^26,28^ The bottomless plate with individual well dimensions matching that of a standard 96 well plate (0.3 cm^2^ culture area) were injection moulded in polystyrene and made in house.

Polystyrene was unsuitable for injection moulding of high aspect ratio nanopillars due to its relatively low glass transition temperature that results in degradation of pillar shapes and mechanical properties. Due to stretching in during injection moulding, use of polycarbonate leads to high aspect ratio nanopillars with features taller and thinner than the quartz master counterparts,^53^ therefore injection moulding of these pillars was carried out using Markrolon®OD2015 Polycabonate. The mechanical properties of the resulting high aspect ratio nanopillars were categorised using finite element modelling (COMSOL Multiphysics) of their morphology under typical Euler-Bernoulli constraints for cantilever beams. Mechanical properties were extrapolated to effective shear and Young’s moduli, using the equations described above.

### Preparation of substrates for cell seeding

All slide arrays were attached to the bottomless multiwell plate by ultrasonic welding (Standard 2000, Rinco Ultrasonics) to create the final multiwell array. Prior to cell seeding, multiwell arrays nanopillars and grooves were cleaned with 70% ethanol and distilled deionized water. then UV sterilized. High aspect ratio pillars were lightly cleaned using compressed air to prevent collapse of nanopillars. All substrates were thereafter treated with O_2_ plasma (80W, 1 minute) then UV sterilized for 20 minutes prior to cell seeding.

### hiPSC-CM cell culture and functionality assays

hiPSC-CM (NCardia) were cultured following the manufacturer’s protocol and proprietary media at a cell density of 100,000 cells/cm^2^. Prior to plating, groove multiwell array were coated with human fibronectin (10 μg/ml, R&D Systems) for 1 hour, then washed twice with phosphate buffered saline (PBS, Sigma-Aldrich). On day 10, cells were loaded with the voltage sensitive fluorescent dye FluoVolt (1:1000, ThermoFisher) along with Powerload (1:100, ThermoFisher) in serum-free medium.and incubated for 25 min at 37°C. Subsequently, action potentials were recorded using the CellOPTIQ^®^ system (Clyde Biosciences) at 10,000 fps as the depolarization time of cardiomyocytes is between 5 and 10 ms. Additionally, contractility analysis was done by recording videos at 100 fps that were analyzed using the MuscleMotion software.^54^

### Chondrocyte cell culture

Isolation of costal chondrocytes were performed as described.^55^ After isolation, murine chondrocytes were cultured in alpha minimum essential medium supplemented with ascorbic acid, glutamate, sodium pyruvate, 10% fetal bovine serum (FBS) and 1% penicillin/streptomycin. Extracted chondrocytes were either used after routine culture in standard tissue culture plastic (cultured chondrocytes) or immediately after harvest (isolated chondrocytes). Chondrocytes were seeded on nanopillars at 2500 cells/cm^2^ in 100 µl complete media, with medium change every 2 days. Chondrocytes were tested for viability, harvested for gene expression analysis at specific timepoints, or fixed for immunohistochemistry and immunofluorescence, as described below.

### MC3T3 cell culture

MC3T3 pre-osteoblasts (ATCC) were cultured using minimum essential medium alpha without ascorbic acid and containing 10% FBS and 1% penicillin/streptomycin for 10 days. MC3T3 were seeded on high aspect ratio nanopillars at 5000 cells/cm^2^ and 100 µl complete media. MC3T3 were harvested for immunofluorescence staining at 24 hours after culture and gene expression analysis after 10 days of culture.

### Immunofluorescence staining and imaging

At selected timepoints, cells were fixed with 4% paraformaldehyde and permeabilized with 0.1% Triton-X 100. Then, samples were blocked with 1% bovine serum albumin and 10% goat serum in PBS. hiPSC-CM were stained against α-actinin (E7732, Sigma, 1:500) using an Alexa Fluor 488-conjugated goat-anti-mouse-antibody (Life Technologies, 1:500) secondary. Chondrocytes and MC3T3 were stained against actin using Alexa Fluor 488-phalloidin (LifeTechnologies). NucBlue fixed cell stain (LifeTechnologies) was used to stain the nuclei of the cells. Imaging was performed under a 10X (numerical aperture 0.3), 20X (numerical aperture 0.45) or 40X magnification (numerical aperture 0.6) using an EVOS FL2 Auto microscope, (ThermoFisher).

### RNA harvest and qPCR

At specified timepoints, total RNA was harvested from cells (ReliaPrep Cell RNA extraction kit, Promega). Relative gene expression was measured from a total of 5 ng RNA using a one-step qPCR kit with SYBR dye (PrimerDesign) and normalized to GAPDH or 18S ribosomal RNA housekeeping gene. A list of the forward and reverse primers used are given in **Table S2**.

### Proliferation rate analysis

Metabolic rate was used as a surrogate marker for chondrocyte proliferation. At selected time points, chondrocytes on nanopillar arrays were added with PrestoBlue reagent (ThermoFisher Scientific, 1:100 dilution). Fluorescence of the reduced reagent was measured at 590 nm emission and 560 nm excitation using a microplate reader (Tecan Infiniti Pro) and was normalized to cultured chondrocytes on Flat at day 7.

### Alcian blue staining and quantification

At different time points, cultured chondrocytes grown on nanopillar arrays were fixed with 4% paraformaldehyde for 15 mins at 4°C. Thereafter, each well was incubated with 0.1% Alcian blue 8GX (Sigma Aldrich) dissolved in 0.1N hydrochloric acid in phosphate buffered saline for 30 mins. Subsequently, a flatbed scanner was used to take color images (at 1200 pixels per image) of the nanopillar arrays. White balance correction of nanopillar array image was performed before image deconvolution to extract the Alcian blue stain. Measurement of Alcian blue intensity was performed using the color deconvolution plugin for ImageJ (National Institutes of Health). All intensity measurements were normalized to those on Flat at day 7.

### Statistical analysis

All data are presented as mean ± standard deviation. Statistical analysis was performed using GraphPad Prism v7.0. One-way ANOVA with Tukey’s post hoc test or two-way ANOVA with Dunnett’s post hoc test was used, with p<0.05 considered significant.

## Author Contributions

NG conceptualized the study. EH and MFAC carried out biological studies on multiwell arrays. FAC, AS, RL and PMR made multiwell arrays. EH, MFAC, FAC, and NG wrote and edited the manuscript with contributions from other authors. All authors have given approval to the final version of the manuscript.

## Funding Sources

European research council FAKIR 648892 Consolidator Award

British Heart Foundation 4-year PhD program

Engineering and Physical Sciences Research Council (EPSRC) PhD funding – ref 1651390

Biotechnology and Biological Sciences Research Council (BBSRC) BB/K011235/1

## Acknowledgements

E Huethorst is supported by the British Heart Foundation 4-year PhD program. MFA Cutiongco is financially supported by the University of Glasgow MG Dunlop Bequest, College of Science and Engineering Scholarship, FAKIR 648892 Consolidator Award from the European Research Council. F Campbell thanks financial support from the EPSRC for his PhD studies. We acknowledge the James Watt Nanofabrication Centre for fabrication work. We thank E Barbour for her assistance with the MC3T3 cell culture. We thank Prof. G Smith for making the hiPSC-CM and CellOPTIQ®system available. We thank Mr. Oliver Sharp from KNT for assistance with the images of fabricated devices.

## Supporting information

Supplementary Tables (Microsoft Word)

Supplementary Videos (avi)

### Abbreviations

ACAN: aggrecan
APD50: action potential duration at 50% of the amplitude
CD50: contraction duration at 50% of the amplitude
CHF_3_/Ar: trifluoromethane / argon
COL1A: collagen type 1a
COL2A: collagen type 2a
DMEM: dulbecco’s modified Eagle’s medium
EBL: electron beam lithography
fps: frames per second
IPA: isopropyl alcohol
MIBK: methyl isobutyl ketone
NiCr: nichrome
NIL: nanoimprint lithography
NMP: N-Methyl-2-pyrrolidone
OCN: osteocalcin
OPN: osteopontin
qPCR: quantitative polymerase chain reaction
RIE: reactive ion etching
RUNX2: Runx family transcription factor 2
SD: standard deviation
SEM: scanning electron microscope
SF_6_ / C_4_F_8_: sulfur hexafluoride / octafluorocyclobutane
SOX9: SRY-box 9
T_contraction_: contraction time
T_relaxation_: relaxation time.

