## Supplementary Tables for "Customizable, engineered substrates for rapid screening of cellular cues"

### Supporting Tables

**Supporting Table 1**. Overview of dimensions and resulting stiffness of high aspect ratio nanopillars.

Effective shear modulus calculated by combining equations (1) and (2):

$$\bar{G}' = \frac{3}{16}\left(\frac{d}{l}\right)^2 fE * \Delta k$$

| **Diameter Tip (nm)** | **Height (µm)** | **Pitch (µm)** | **Fill Factor** | **Individual pillar spring constant (k, pN/nm)** | **Poisson ratio (ν)** | **Effective Shear Modulus ($\bar{G}'$, kPa)** |
|---|---|---|---|---|---|---|
| 70 | 2.35 | 1 | 0.0044 | 3.9 | 0.37 | 9 |
| 70 | 1.6 | 1 | 0.0044 | 10.6 | 0.37 | 16 |
| 90 | 1.35 | 1 | 0.0064 | 51.7 | 0.37 | 70 |
| 100 | 1.9 | 0.5 | 0.0314 | 11.4 | 0.37 | 87 |
| 115 | 1.95 | 1 | 0.0104 | 47.8 | 0.37 | 93 |

**Supplementary Table 2.** Primers used for measuring gene expression of cardiomyogenic, chondrogenic and osteogenic function.

| Mouse gene name | | Primer Sequence | Amplicon size |
|---|---|---|---|
| 18s ribosomal RNA (housekeeping gene) | Fwd<br>Rev | AAGTCCCTGCCCTTTGTACACA<br>GATCCGAGGGCCTCACTAAAC | 100 |
| *COL1A1* (Collagen type 1 alpha 1 chain) | Fwd<br>Rev | AACGAGATCGAGCTCAGAGG<br>GACTGTCTTGCCCCAAGTTC | 99 |
| *COL2A1* (Collagen type 2 alpha 1 chain) | Fwd<br>Rev | ACGAAGCGGCTGGCAACCTCA<br>CCCTCGGCCCTCATCTCTACATCA | 73 |
| *COL10A1* (Collagen type 10 alpha 1 chain) | Fwd<br>Rev | TTCTCCTACCACGTGCATGTG<br>AGGCCGTTTGATTCTGCATT | 191 |
| *ACAN* (Aggrecan) | Fwd<br>Rev | GTGAGGACCTGGTAGTGCGAGTGA<br>GAGCCTGGGCGATAGTGGAATATA | 103 |
| *SOX9* (SRY-Box 9) | Fwd<br>Rev | GGAAGGGAGAGAGAGAGAGAAA<br>CGGGATTTAAGGCTCAAGGT | 138 |
| *RUNX2* (Runx family transcription factor 2) | Fwd<br>Rev | AAGTGCGGTGCAAACTTTCT<br>TCTCGGTGGCTGCTAGTGA | 90 |
| *OCN* (Osteocalcin) | Fwd<br>Rev | CTGACCTCACAGATGCCAAG<br>GTAGCGCCGGAGTCTGTTC | 98 |
| *OPN* (Osteopontin) | Fwd<br>Rev | TCAGGACAACAACGGAAAGGG<br>GGAACTTGCTTGACTATCGATCAC | 139 |
| *MYOG* (Myogenin) | Fwd<br>Rev | GAGACATCCCCCTATTTCTACCA<br>GCTCAGTCCGCTCATAGCC | 106 |
| *MYH7* (Myosin heavy chain 7) | Fwd<br>Rev | CTCAAGCTGCTCAGCAATCTATTT<br>GGAGCGCAAGTTTGTCATAAGT | 153 |
| *ASF1B* (Anti-silencing function 1B histone chaperone) | Fwd<br>Rev | CCTTCCGGTTCGAGATCAGC<br>GGATGGGTTTGGGGCATCAG | 185 |
